# Light spectra modify nitrogen assimilation and nitrogen content in *Quercus variabilis* Blume seedling components: A bioassay with ^15^N pulses

**DOI:** 10.1101/2020.12.02.407924

**Authors:** Jun Gao, Jinsong Zhang, Chunxia He, Qirui Wang

## Abstract

The light spectra that reach plants change across different shading conditions, may alter the pattern of nitrogen (N) uptake and assimilation by understory regenerations that are also exposed to N deposition. We conducted a bioassay on Chinese cork oak (*Quercus variabilis* Blume) seedlings subjected to five-month N pulsing with ^15^NH_4_Cl (10.39 atom %) at 120 mg ^15^N plant^-1^ under the blue (48.5% blue, 33.7% green, and 17.8% red), red (14.6% blue, 71.7% red, 13.7% green), and green (17.4% blue, 26.2% red, 56.4% green) spectra provided by light-emitting diodes (LEDs). Half of the seedlings were fed twice a week using a 250 ppm N solution with added phosphorus, potassium, and micro-nutrients, while the other half received only distilled water. Neither treatment affected growth of height, diameter, or leaf area. Compared to the red light spectrum, the blue light treatment increased chlorophyll and soluble protein contents and glutamine synthetase (GS) activity, root N concentration, and N derived from the pulses. The green light spectrum induced more biomass to allocate to the roots and a higher percentage of N derived from internal reserves compared to the other two spectra. The ^15^N pulses demonstrated no interaction with spectra but weakened the reliance on N remobilization from acorns, strengthened biomass allocation to shoots, and induced higher chlorophyll content, GS activity, and N concentration. In conclusion, the red light spectrum should be avoided for *Q. variabilis* regenerations whose biomass allocation to underground organs are weakened under this condition.

## Introduction

Light is the most important among all environmental factors for plant growth as a determinant of photosynthesis. Early studies on the photoperiodism of juvenile trees were mainly conducted by implementing fluorescent and sodium vapor lamps [1–3]. With the development of high-intensity discharge illuminators, high-pressure sodium (HPS) lamps were found to emit light that can more efficiently benefit plants compared to traditional lighting sources [4–6]. In the last decade, the development of light-emitting diodes (LEDs) broadened the range of choices for light sources in tree seedling cultures [7]. Many studies have demonstrated the advantages of continuous LED lighting over HPS or fluorescent sources for culturing high quality tree seedlings [5, 8–16]. Light from LED sources has been established to be a solid and flexible instrument for current tests on the response of juvenile trees to different spectra.

The various responses of woody plants to different light spectra result from the natural selection to acclimate to changes in light qualities in the understory layer [17]. In studies on tree seedlings under varied spectra conditions created by LED, the effects on mass production and physiology were determined using growth and biomass parameters [5, 8, 15, 16], photosynthesis and stomatal conductance [5, 8, 16], and transplant performance [8, 15]. All current studies were conducted using the horticultural system of prolonged photoperiod via continuous lighting [8, 9, 14] or supplemental illumination [5, 15, 16]. However, experts found in the 1960s that a prolonged photoperiod can generate an interaction with fertilizer regime on tree seedlings [18]. This interaction was further tested by Wei et al. [4], and a negative effect from longer-photoperiod caused by nutrient dilution has been reported [4, 8], which was suggested by the classic model of the relationship between nutritional status and growth pattern in tree seedlings [19]. In more recent studies, the emphasis of the effect of spectra on tree seedlings switched to detecting the effects of spectra combined with fertilizer regimes [10, 13].

Studies revealed that the HPS spectrum in continuous lighting results in remarkable nutrition dilution in the absence of a proper regime of additional nutrient input [4, 8]. Nutrient dilution in a tree seedling is a natural response to acceleration of biomass accumulation without sufficient nutrient supply at the late stage of seedling growth [20]. Nutrient dilution intensively impairs seedling quality and negatively influences outplant performance [20, 21]. An LED spectrum with a higher proportion of red light (600–700 nm) would increase nutrition dilution more than HPS light sources [9, 14]. The high blue light spectrum (400–500 nm) was found to accelerate biomass accumulation, which stimulated the occurrence of nutrient dilution [15, 16]. The white-light spectrum with a high green light content (500-600 nm) could also induce nutrient dilution. As shown by Zhao et al. [10] and Wei et al. [13], a higher rate of fertilizer supply can generally solve the issue of nutrient insufficiency and counter nutrient dilution in LED-lighted seedlings. As nutrient content can be supplied by either exogeneous fertilizers or endogenous reserves [22–25], further explanation of the percent and amount of inner-derived nutrients by exposure to LED spectra is lacking.

Under global climate change, anthropogenic activities have caused the increase in atmosphere nitrogen (N) deposition [26–28]. Intensive N input to forest ecosystems has altered the C and N cycling in tree populations [29]. The combined effect of sunlight spectra and N deposition is influencing understory regenerations in forest communities, which, however, has attracted surprisingly little attention. The most frequent employment of photosynthetic photon flux density (PPFD) at tree-perceived level in the above-mentioned LED lighting experiments ranged from 60–80 μmol m^-2^ s^-1^ [5, 9], which fully falls in the range of understory sunlight transmittance [17]. The results of the above-mentioned studies demonstrate a knowledge gap that can be easily filled by new observations with studies using LED and N pulse to mimic natural conditions.

Chinese cork oak (*Quercus variabilis* Blume) is a widely distributed deciduous broadleaf tree species in East Asia across temperate and subtropical regions (24° to 42° N and 96° to 140° E) [30, 31]. This oak is a valued hardwood species that can be used as the raw material for construction, furniture, and nortriterpenoid extracting [32]. Because of the high efficiency of N resorption but low growing speed [30], Chinese cork oak spends a long time as regenerated saplings in the understory layer, where it is more sensitive to N deposition than other oak species [33]. The frequent habit to remobilize N acorns not only increases the reliance on inner-derived N in acorns to feed current growth but also increases the complexity of distinguishing different sources of N in total content [34]. In this study, Chinese cork oak was raised in a bioassay under controlled conditions with different LED light spectra and simulated N deposition. The exogenous N input was labeled using a ^15^N-isotope to facilitate distinguishing different sources of derived N. We hypothesized that: (1) the red light spectrum would induce more N derived from the pulse (NDFP) than in blue and green lights due to promoting vigorous growth, (2) green light would promote N derived from reserves (NDFR) to compensate for reduced demand, and (3) NDFR in acorns would be benefited by green light.

## Materials and Methods

### Plant Material and Growing Condition

Pre-germinated Chinese cork oak acorns were collected from a seed source in Jigongshan National Natural Reserve (31°46’–31°52’ N, 114°01’-114°06’ E) in central China. Acorns were sterilized in potassium permanganate solution (0.5%, w/w) for 30 min and sown into 212 cm^3^ (7 × 4 × 13 cm, top diameter × bottom diameter × height) volume trayed-cavities (4 × 8 individuals embedded in a tray) filled with N-poor substrates (Mashiro-Dust^™^, Zhiluntuowei A&F S&T Inc., Changchun, China). This growing medium was comprised of 70% peat, 10% spent mushroom residue (SMR), and 20% perlite. Chemical analysis revealed that this media contained 9.8 mg kg^-1^ NH_4_–N, 5.6 mg kg^-1^ NO_3_-N, 870.3 mg kg^-1^ PO_4_-P, pH of 5.3, and an electrical conductance (EC) of 0.86 dS m^-1^. Trayed acorns were incubated in the Laboratory of Combined Manipulations of Illumination and Fertility on Plant Growth (Zhilunpudao Agric. S&T Inc., Changchun, China) for 51 days until 80% germination. During germination, the indoor environment was controlled to 29/21 °C (day/night) and relative humidity to 58/61% (max/min). Germinated seedlings with fine roots attached to the acorn were transplanted to pots (11.5 × 7.5 × 9.5 cm, top diameter × bottom diameter × height) with one individual in each pot. Pots were filled with the same growing medium as described above. Four pots were placed in one tank (40 × 60 cm, width × length). Finally, a total of 216 potted seedlings were placed in 54 tanks.

### LED Lighting Treatment

The tanked pots of seedlings were placed on iron shelves (2.0 × 0.5 ×1.5 m, height × width × length). Each shelf had three floors of spaces as growing chambers (0.5 × 0.5 × 1.5 m, height × width × length). Two tanks were placed in one chamber, so each shelf contained six tanks with 24 seedlings. An LED panel (0.5 × 1.2 m, width × length; Pudao Photoelectricity, Zhiluntuowei A&F S&T., Inc., Changchun, China) was attached to the ceiling of each chamber. The LED panel provided lighting in a n18 h photoperiod, which has been proven to benefit the growth of slowly-growing species [8, 35]. One hundred diodes were embedded in the downward surface of panel at a spacing of 2 × 2 cm. Every diode was designed to emit one of blue, red, or green light. The electric flow for diodes emitting the same type of light was controlled by one electrical administration transformer. Electric power for the array of red-light diodes was administrated by a 200 W transformer, while power for arrays of either green- or blue-lights diodes was administrated by a 135 W transformer. Changing the electric flow for one array of diodes can modify the photosynthetic photon flux rate (PPFD) intensity for the color of lights. Therefore, the spectrum of light that was emitted by one LED-panel was obtained by adjusting the electric flow for the three types of arrayed diodes. The visual color of light from a LED panel is a mixture of wavelengths for blue, red, and green lights at various PPFD intensities that were controlled by the electric flow.

Visually blue, red, and green lights were employed as mixed-wavelengths in visible lights from 400 to 700 nm (EVERFINE PLA-20, Yuanfang Elect. S&T Inc., Hangzhou, China) [36]. As our aim was to test the effect of different spectra, the intensity of PPFD was controlled to a similar level that produces ordinary growth [5, 15, 16] to eliminate the unexpected error from the interplay between changes in both lighting spectra and intensity [9, 10, 12–14]. The spectra for three visual colors of lights that fulfilled the above requirements were generated as follows:

i. Blue light spectrum: Electric flow was adjusted to 70%, 10%, and 10% of full power for arrayed diodes emitting blue, red, and green lights, respectively.
ii. Red light spectrum: Electric flow was adjusted to 10%, 30%, and 20% of full power for arrayed diodes emitting blue, red, and green lights, respectively.
iii. Green light spectrum: Electric flow was adjusted to 30%, 20%, and 100% of full power for arrayed diodes emitting blue, red, and green lights, respectively.

The outcome of the three spectra is shown in Figure 1 and the specific optical characteristics for the three spectra are provided in Table 1.

**Figure 1.**
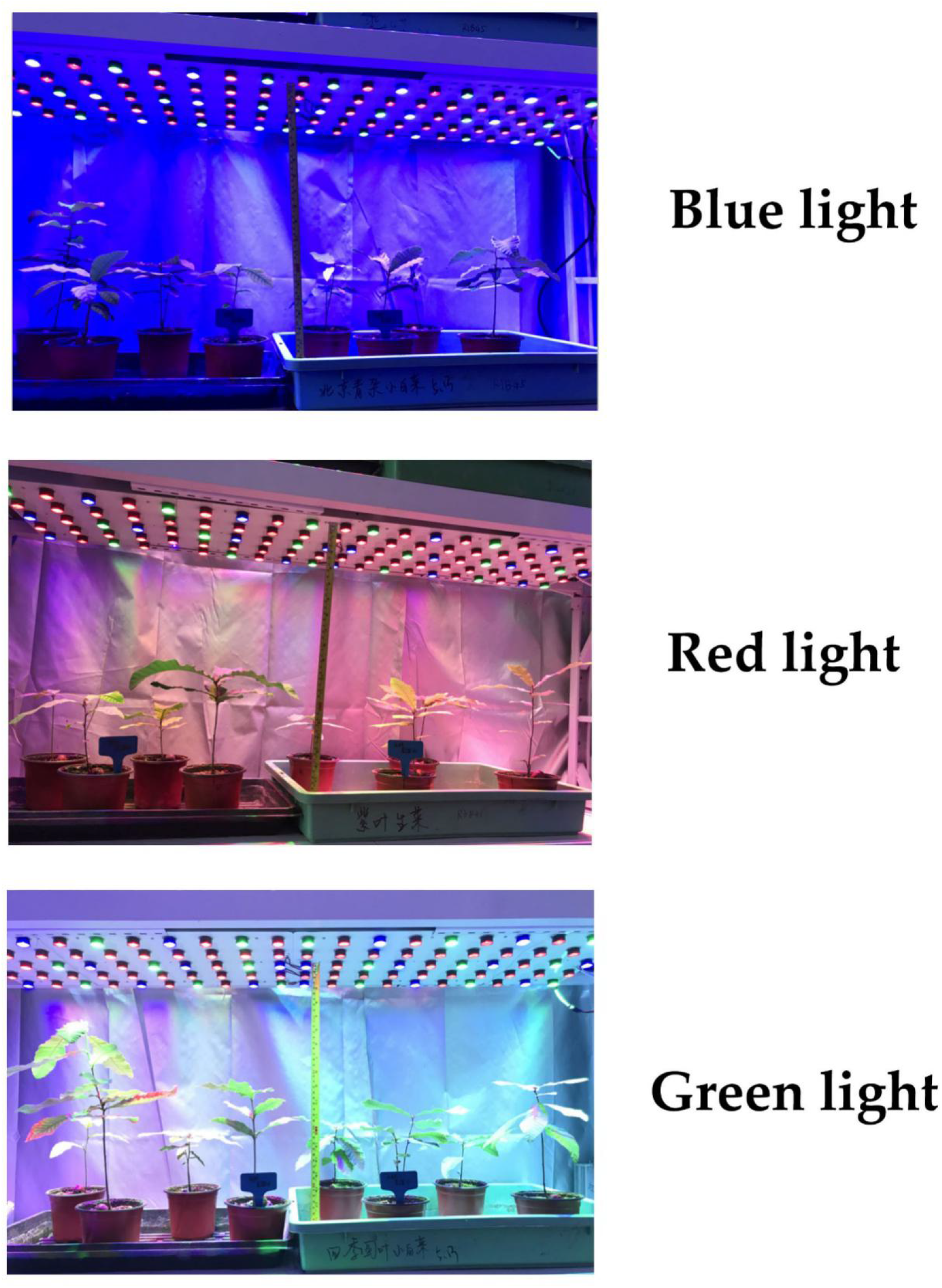
Typical performance of Chinese cork oak (*Quercus variabilis* Blume) seedlings exposed to blue (48.5% blue, 33.7% green, and 17.8% red), red (14.6% blue, 71.7% red, and 13.7% green), and green (17.4% blue, 26.2% red, 56.4% green) colors of light-emitting diode (LED) spectra. Tanks for potted seedlings were 40 × 60 cm. Green tanks contained seedlings subjected to ^15^N pulses and black tanks contained seedlings subjected to water addition of the same volume.

**Table 1.** Optical characteristics (40 cm beneath the lighting source) of blue, green, and red visible LED lighting of combined wavelengths in a wide bandwidth from 400 to 700 nm.

| Optical characteristics |  | Visible light color |  |  |
| --- | --- | --- | --- | --- |
|  |  | Blue | Red | Green |
| PPFD ($\mu\text{mol m}^{-2} \text{s}^{-1}$) | | 95.18 | 97.86 | 96.46 |
| Proportional percent | Blue | 17.8% | 14.6% | 17.4% |
|  | Red | 33.7% | 71.7% | 26.2% |
|  | Green | 48.5% | 13.7% | 56.4% |
|  | Total | 100% | 100% | 100% |

Note: PPFD, photosynthetic photon flux density.

### ^15^N Pulsing Treatment

In this study, N was delivered to Chinese cork oak seedlings through pulsing at a rate of 19.6 kg N ha^-1^ (~120 mg N plant^-1^), which is the annual amount of wet N deposition for the natural population in the region of Jigongshan National Natural Reserve [37]. This amount is equal to 117 mg N plant^-1^, as four individuals were arranged in a 0.24 m^2^ area, using the N consumption of mature trees to pulse our seedlings. It was reported that 125 mg N plant^-1^ can maximize dry mass production at the sufficiency level of N input for oak (*Quercus ilex*) seedlings [38]. N was pulsed according to the method in previous studies [39, 40]. Seedlings were fed twice a week using a balanced nutrient solution [23] with 250 ppm N delivered through ammonium chloride (NH_4_Cl). Briefly, the solution contained 60 ppm phosphorus and 100 ppm potassium with micro-nutrients added [23]. Exogenous N was labeled by ^15^N as 10.39 atom % ^15^NH_4_Cl (Shanghai Research Institute of Chemical Industry, Yunling Road, Shanghai, China). A 10 mL nutrient solution was applied to the soil surface of each pot using a 5 mL pipettor and only soils were pulsed to avoid ^15^N contamination of shoots [39, 40]. N was pulsed through 40 applications in five months. Only one of the two tanked pots of seedlings received ^15^N pulsing, while the other group of seedlings received distilled water at the same volume as the control.

### Experiment Design

This study was conducted as a split-block design with the main block as three LED spectra, each of which harbored two ^15^N pulse treatments. Three iron shelves with LED panels emitting blue, red, and green lights were assigned to one block, and three blocks of shelves were assigned as three replicates that were randomly placed. Four seedlings in one tank (either ^15^N pulse or not) per shelf level were assigned as the basic unit of sampling and measurement. Three levels of tanked pots of seedlings were grouped to average the observations for combined spectrum and N treatment.

### Sampling and Measurement

All four seedlings per tank were sampled and measured for height and root collar diameter (RCD). Four seedlings were assigned to two groups, with two randomly chosen seedlings per group. One group of seedlings were separated into shoots (leaves and woody stems), roots, and acorn. Roots were washed three times, by tap water once and distilled water twice, to carefully remove substrates without causing damage to the fine roots. All three parts of the seedlings for each group were dried in oven at 65 °C to constant mass then weighed, ground, and measured for total N concentration and ^15^N enrichment using a stable isotope ratio mass spectrometer (Finnigan DELTA^plus^ XP, Thermo Fisher Scientific, Grand Island, NY, USA) [41]. N content was calculated by the product of N concentration and biomass. Leaves of the other group of seedlings were directly used for determination or excised and stored in tinfoil-folded liquid N until measured for physiological parameters.

Four leaves were randomly chosen from the two seedlings and scanned to obtain a 300 dots per inch (dpi) image (HP Deskjet 1510 scanner, HP Inc., Palo Alto, CA, USA), whose background was stratified and removed in Photoshop CS V. 8.0 (Adobe, San Jose, CA, USA). The front layer of leaves was opened as a histogram. The leaf green index (GI) can be directly read from the background data of the histogram [42–44]. Leaf area can be calculated as the total pixels of the histogram divided by the square of the dpi [43–45]. Scanned leaves were oven-dried at 65 °C to constant weight and measured for the biomass of a single leaf. Thereafter, the specific leaf area (SLA) was calculated as the product of leaf area and single leaf biomass.

Chlorophyll and protein concentrations were analyzed on fresh leaves using the method adapted from Zhao et al. [10]. Briefly, chlorophyll was determined using a 0.05 g sample, which was placed in a hydraulic bath at 65 °C for 1 h and used to determine concentrations of chlorophyll-a, chlorophyll-b, and carotenoid at wavelengths of 663, 645, and 470 nm, respectively. Protein concentration was measured on a 0.1 g sample that was ground in 1 mL of phosphate buffer at pH 7.5, centrifuged at 3000 rpm for 10 min, treated with 0.1 mL of Folin’s reagent, and determined at 650 nm.

Foliar glutamine synthetase (GS) activity was assessed using the method reported by Wei et al. [45]. A 0.5 g leaf sample was homogenized in 5 mL extraction buffer (3.059 g Tris, 0.249 g MgSO_4_·7H_2_O, 0.309 g dithiothreitol, and 68.5 g sucrose dissolved in 500 mL deionized water brought to pH 8.0 using 0.05 mM HCl) at 6000 rpm for 20 min. We added 0.7 mL supernatant to 6 mL reaction B (6.118 g Tris, 9.959 g MgSO_4_·7H_2_O, 1.726 g monosodium glutamate, 1.211 g cysteine, 0.192 g Triethylene glycol diamine tetraacetic acid (EGTA), pH 7.4, 500 mL) with the solution of 0.7 mL ATP (40 mM). The reaction mixture was incubated at 37 °C for 30 min and stopped by adding 1.0 mL of ferric chloride reagent (3.317 g trichloroacetic acid, 10.102 g FeCl_3_·6H_2_O, 5 mL sulfuric acid, 100 mL). The absorbance of the product, glutamyl-γ-hydroxamate, was measured at 540 nm. Protein content was determined using the Folin’ reagent method as described above.

### Calculations

We calculated the isotopic abundance for N in atom % (*A*_N_%) as [40, 46]:

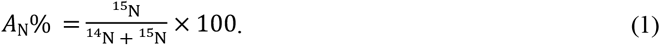

The NDFP percentage was calculated as the relative specific allocation of pulse N (*RSA*_N_%) as [25, 40, 47]:

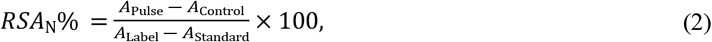

where *A*_Pulse_ is the ^15^N abundance in pulsed seedling organs (shoots, roots, and acorns), *A*_Control_ is the ^15^N abundance in organs of controlled seedlings without pulsing, *A*_Labd_ is the ^15^N abundance in (^15^NH_4_)_2_Cl solution, and *A*_Standard_ is the ambient standard of 0.366% [25, 47]. The NDFP amount can be calculated by:

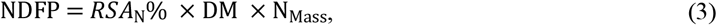

where DM is the dry mass of the seedling organ and N_Mass_ is the N concentration in this organ. Therefore, the NDFR percentage can be calculated as 100% minus *RSA*_N_% [40]. The NDFR amount can be calculated as [40]:

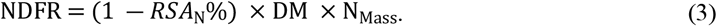

### Statistical Analysis

Data were analyzed using SAS (ver. 9.4 64-bit, SAS Institute Inc., Cary, NC, USA). Shapiro–Wilk test was conducted on data using the univariate procedure, and logarithm transformation plus 1.0 was used for data with abnormal distribution to meet the normality requirement [48]. The effects of LED spectra (blue, red, and green) and ^15^N pulse (pulse vs control) were tested using two-way analysis of variance (ANOVA) using a split-block model with the placement of three replicated blocks (*n* = 3) as the random factor in the mixed procedure. When a significant effect was indicated under the condition of over-95% probability with degrees of freedom for the model and error of 5 and 12, respectively, means were arranged and compared according to Tukey’s test at the 0.05 level.

## Results

### Growth and Leaf Morphology

Neither LED spectra nor ^15^N pulse produced a significant effect on height or RCD (Table 2). Seedling height ranged between 16.70 ± 2.93 and 19.13 ± 1.91 cm, and RCD ranged from 0.35 ± 0.02–0.42 ± 0.05 cm in response to combined spectra and ^15^N pulse treatments. The two treatments produced no effect on LA or GI (Table 2), which ranged from 83.36 ± 7.03–107.55 ± 11.20 cm^2^ and 86.49 ± 1.25–100.29 ± 16.92, respectively. The LED spectra treatment showed a significant effect on SLA (Table 2), which was lower in the red light spectrum (512.84 ± 38.27 cm^2^ g^-1^) than in the blue (440.29 ± 45.50 cm^2^ g^-1^) and green (395.40 ± 21.83 cm^2^ g^-1^) light spectra.

**Table 2.**
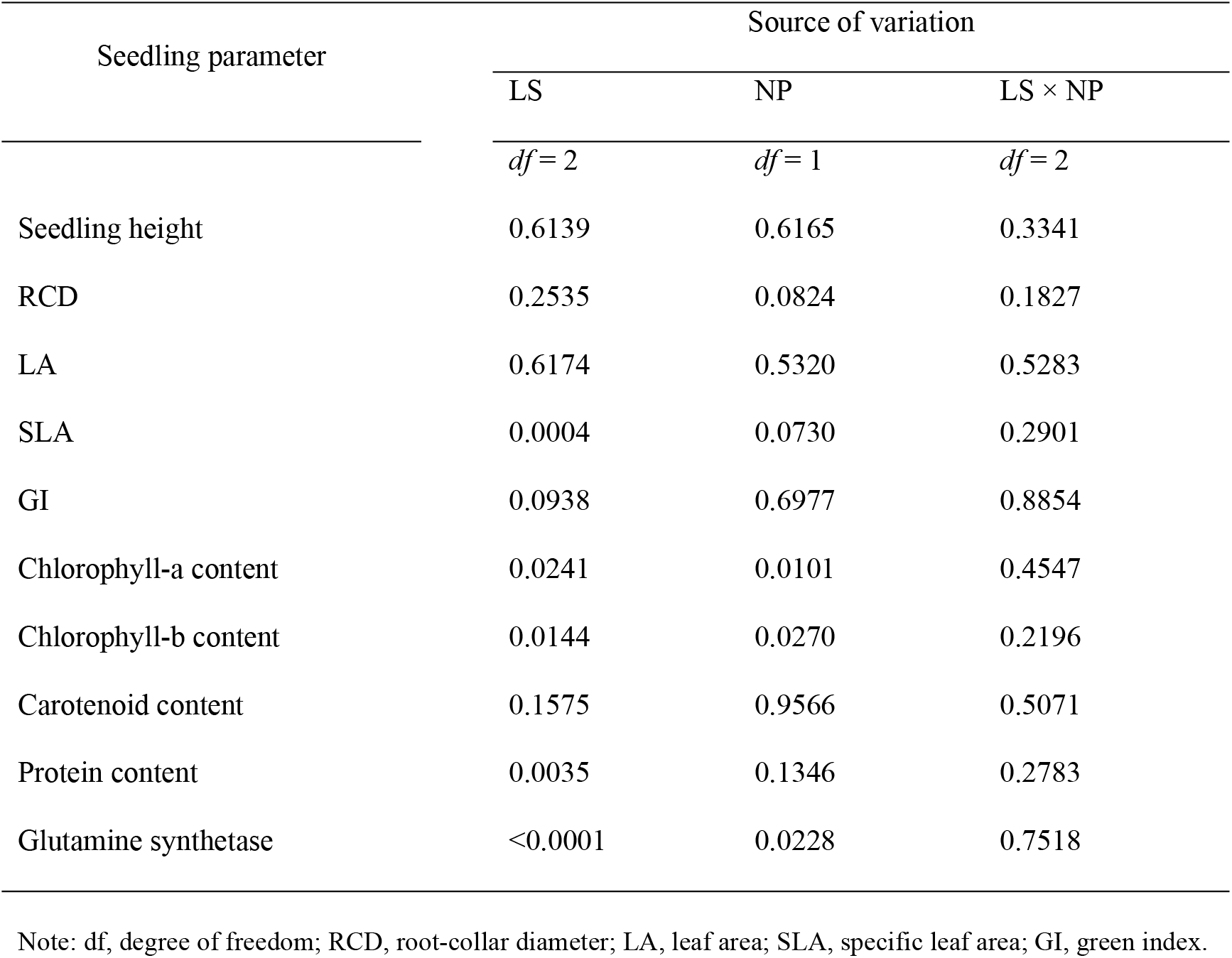
*P*-values from analysis of variance (ANOVA) of lighting spectrum (LS), ^15^N pulse (NP), and their interaction (LS × NP) on growth and foliar physiology in Chinese cork oak (*Quercus variabilis* Blume) seedlings.

| Seedling parameter | Source of variation |  |  |
| --- | --- | --- | --- |
| | LS | NP | LS × NP |
|  | *df* = 2 | *df* = 1 | *df* = 2 |
| Seedling height | 0.6139 | 0.6165 | 0.3341 |
| RCD | 0.2535 | 0.0824 | 0.1827 |
| LA | 0.6174 | 0.5320 | 0.5283 |
| SLA | 0.0004 | 0.0730 | 0.2901 |
| GI | 0.0938 | 0.6977 | 0.8854 |
| Chlorophyll-a content | 0.0241 | 0.0101 | 0.4547 |
| Chlorophyll-b content | 0.0144 | 0.0270 | 0.2196 |
| Carotenoid content | 0.1575 | 0.9566 | 0.5071 |
| Protein content | 0.0035 | 0.1346 | 0.2783 |
| Glutamine synthetase | <0.0001 | 0.0228 | 0.7518 |

Note: *df*, degree of freedom; RCD, root-collar diameter; LA, leaf area; SLA, specific leaf area; GI, green index.

### Leaf Physiology

Either lighting spectra or ^15^N pulse had a significant effect on chlorophyll-a and chlorophyll-b contents (Table 2). Both chlorophyll-a and chlorophyll-b contents were higher in the blue light spectrum than in the red light spectrum (Figure 2A,B). The ^15^N pulse increased the contents of both chlorophyll-a and chlorophyll-b relative to the control (Figure 2D,E). No effect was found on carotenoid content, which ranged between 0.15 and 0.25 mg g^-1^ (Figure 2C,F).

**Figure 2.**
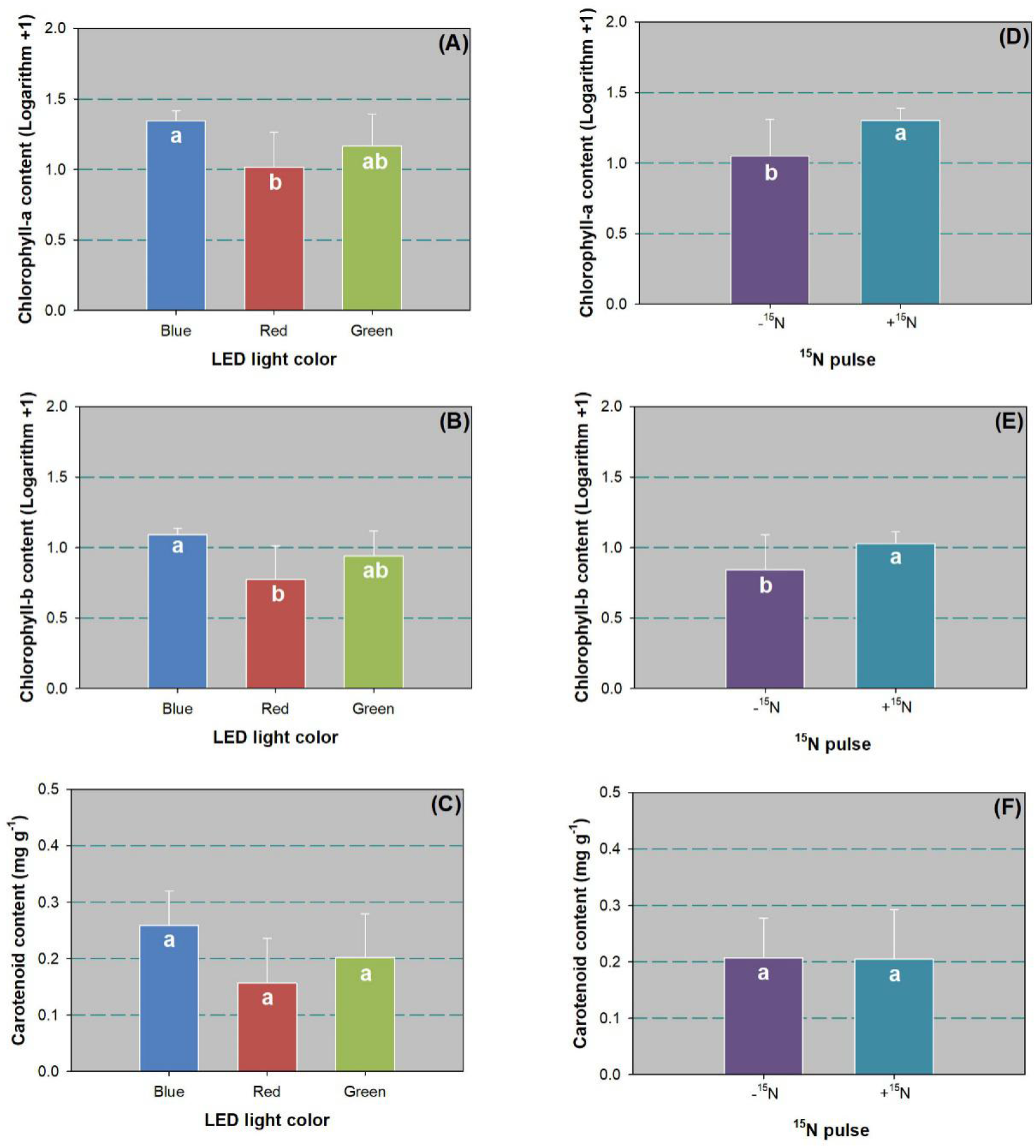
Contents of chlorophyll-a (A,D), chlorophyll-b (B,E), and carotenoid (C,F) in leaves of Chinese cork oak (*Quercus variabilis* Blume) seedlings subjected to blue, red, and green LED spectra (*n* = 24 seedlings per replicate) or subjected to N pulse at 120 mg ^15^N plant^-1^ (+^15^N) or zero (-^15^N) (*n* = 36). Data that failed to follow a normal distribution were transformed by logarithm plus 1.0. Columns and bars represent means and standard errors, respectively. Different letters denote a significant difference according to Tukey’s test at the 0.05 level.

The LED spectra had a significant effect on leaf protein content (Table 2), which was higher in the blue light spectrum than spectra from red and green lights (Figure 3A). The ^15^N pulse did not have any impact on leaf protein content (Figure 3C). Both LED spectra and ^15^N pulses had a significant effect on GS activity (Table 2). Again, the blue light spectrum resulted in higher GS activity than red and green spectra (Figure 3B). The ^15^N pulse also produced a significant increase in GS activity compared to the control (Figure 3D).

**Figure 3.**
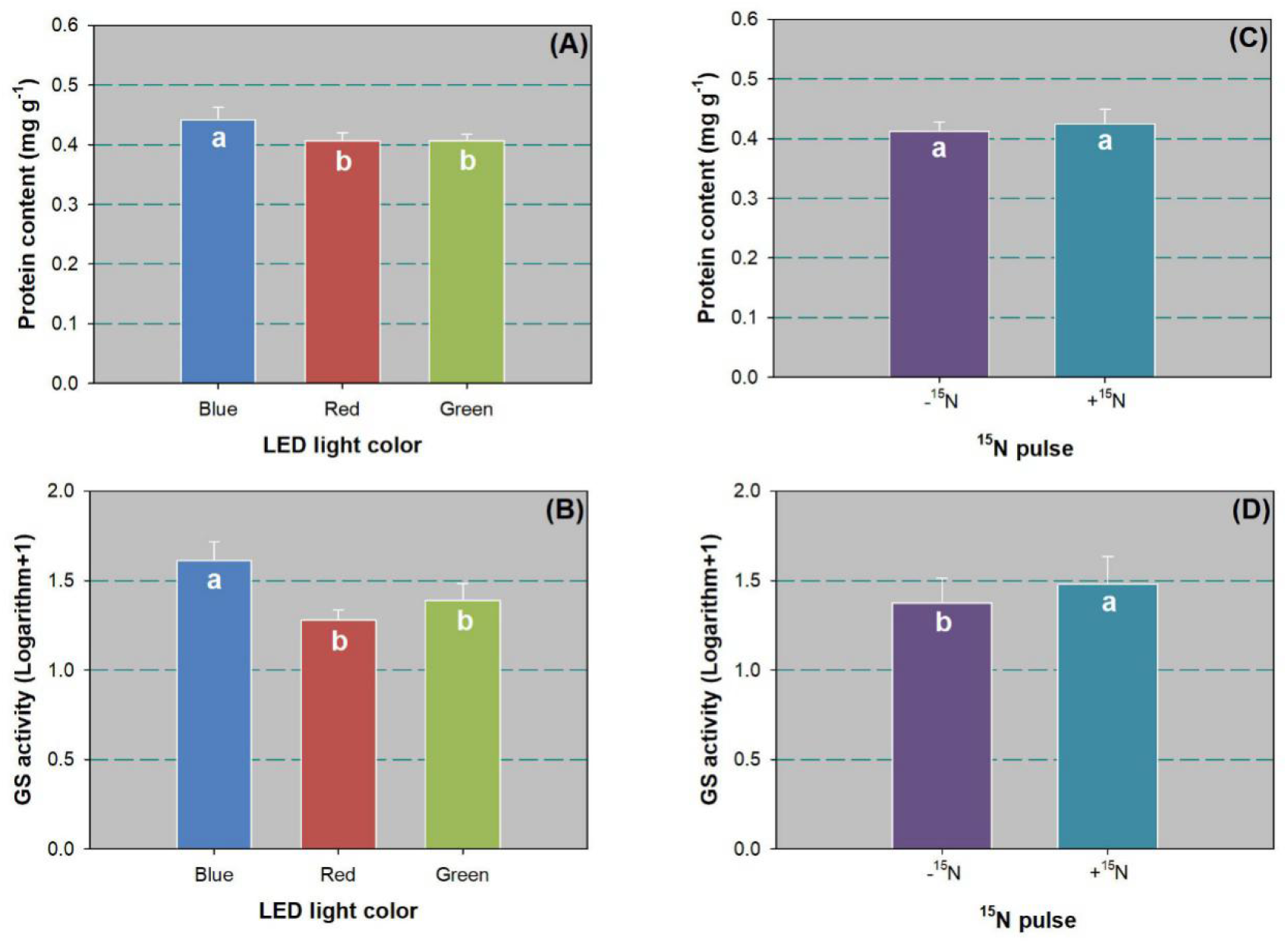
Soluble protein content (A,C) and glutamine synthetase (GS) activity (B,D) in leaves of Chinese cork oak (*Quercus variabilis* Blume) seedlings subjected to blue, red, and green LED spectra lighting (*n* = 24) or subjected to N pulse at 120 mg ^15^N plant^-1^ (+^15^N) or zero (-^15^N) (*n* = 36). Data that failed to follow a normal distribution were transformed by logarithm plus 1.0. Columns and bars represent means and standard errors, respectively. Different letters mark significant difference according to Tukey’s test at the 0.05 level.

### Biomass Accumulation and Allocation

Both LED spectra and ^15^N pulse had a significant effect on biomass in shoot and root parts (Table 3). The green light spectrum led to lower shoot biomass than the blue and red spectra (Figure 4A). In contrast, root biomass in the green light spectrum was higher than in the other two spectra (Figure 4B). Compared with the control, the ^15^N pulse treatment resulted in larger shoot biomass (Figure D), but root biomass was lower (Figure 4E). The ^15^N pulse also lowered acorn biomass relative to the control (Figure 4F).

**Table 3.** *P*-values from ANOVA of lighting spectrum (LS), ^15^N pulse (NP), and their interaction (LS × NP) on biomass and nitrogen (N) concentration and content in aboveground and belowground parts of Chinese cork oak (*Quercus variabilis* Blume) seedlings.

| Seedling parameter | Source of variation |  |  |
| --- | --- | --- | --- |
|  | LS | NP | LS × NP |
|  | *df* = 2 | *df* = 1 | *df* = 2 |
| Shoot biomass | 0.0067 | 0.0013 | 0.8486 |
| Root biomass | 0.0085 | 0.0037 | 0.9492 |
| Acorn biomass | 0.6218 | 0.0225 | 0.2192 |
| Shoot N concentration | 0.0164 | 0.0023 | 0.6068 |
| Root N concentration | 0.0023 | 0.0312 | 0.2200 |
| Acorn N concentration | 0.2118 | 0.0046 | 0.5194 |
| Shoot N content | 0.4390 | 0.0004 | 0.8771 |
| Root N content | 0.0020 | 0.8409 | 0.4896 |
| Acorn N content | 0.0134 | 0.0247 | 0.1168 |

**Figure 4.**
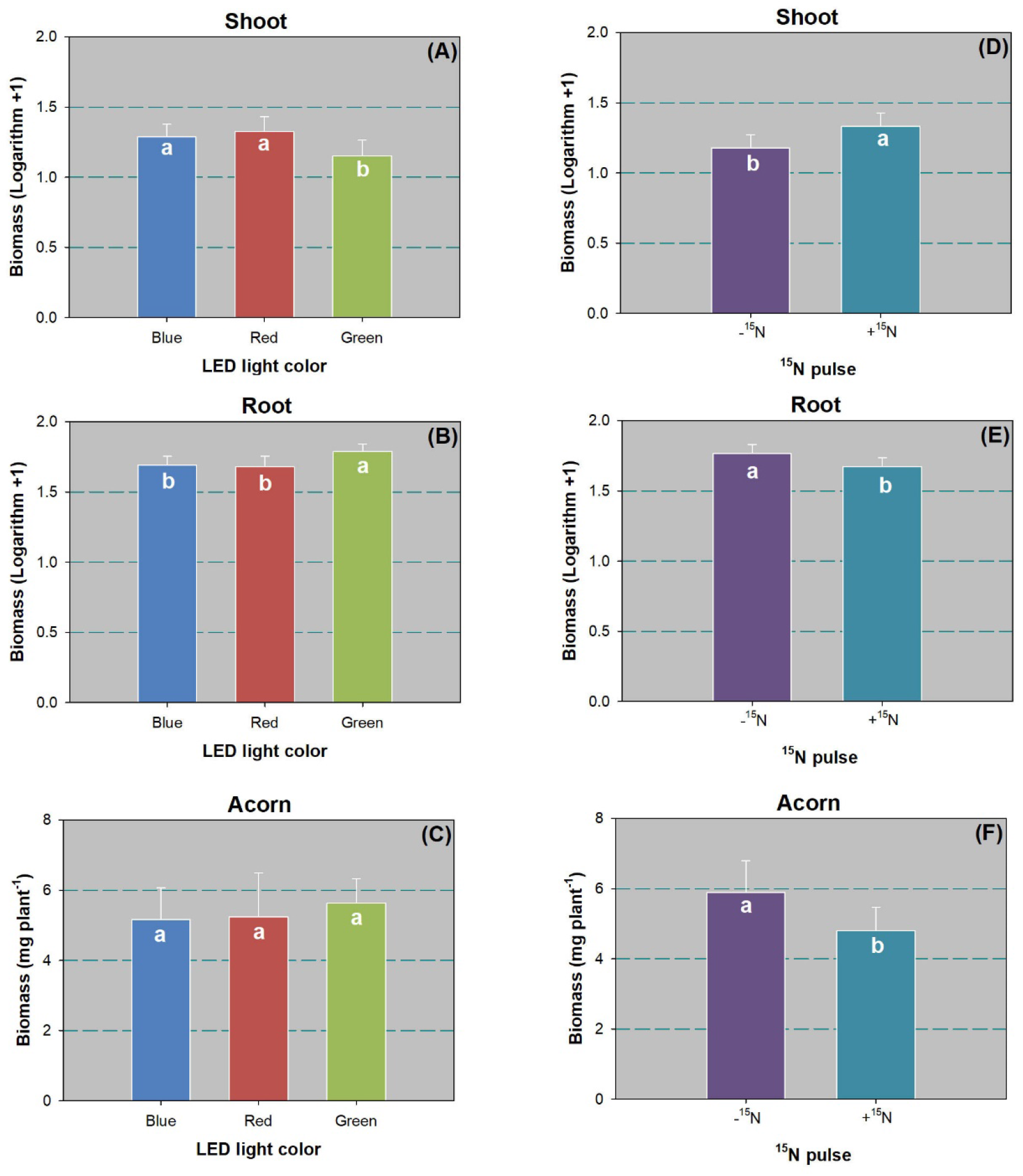
Biomass in shoot (A,D), root (B,E), and acorn (C,F) of Chinese cork oak (*Quercus variabilis* Blume) seedlings subjected to blue, red, and green colors of LED spectra (*n* = 24) or subjected to N pulse at 120 mg ^15^N plant^-1^ (+^15^N) or zero (-^15^N) (*n* = 36). Data that failed to follow a normal distribution were transformed by logarithm plus 1.0. Columns and bars represent means and standard errors, respectively. Different letters denote a significant difference according to Tukey’s test at the 0.05 level.

Both LED spectra and ^15^N pulse treatments had a significant effect on root to shoot biomass ratio (R/S) (F_5,12_ = 21.51; *P* < 0.0001). R/S was higher in the green light spectrum (5.62 ± 1.66) than in the blue (3.25 ± 0.73) and red (3.06 ± 1.07) spectra. The ^15^N pulse lowered R/S by 41% relative to the control (R/S values: 2.97 ± 0.89 and 4.99 ± 1.57, respectively).

### N Concentration

Both LED spectra and ^15^N pulse treatments had a significant effect on N concentration in most seedling tissue parts except for acorn (Table 3). Shoot N concentration was higher in the green light spectrum than in the red light spectrum (Figure 5A). However, root N concentration was higher in the blue light spectrum than in the red light spectrum (Figure 5B). The ^15^N pulse resulted in higher N concentration in both shoots (Figure 5E) and roots (Figure 5F) compared to the control. In contrast, the ^15^N pulse lowered acorn N concentration relative to the control (Figure 5F).

**Figure 5.**
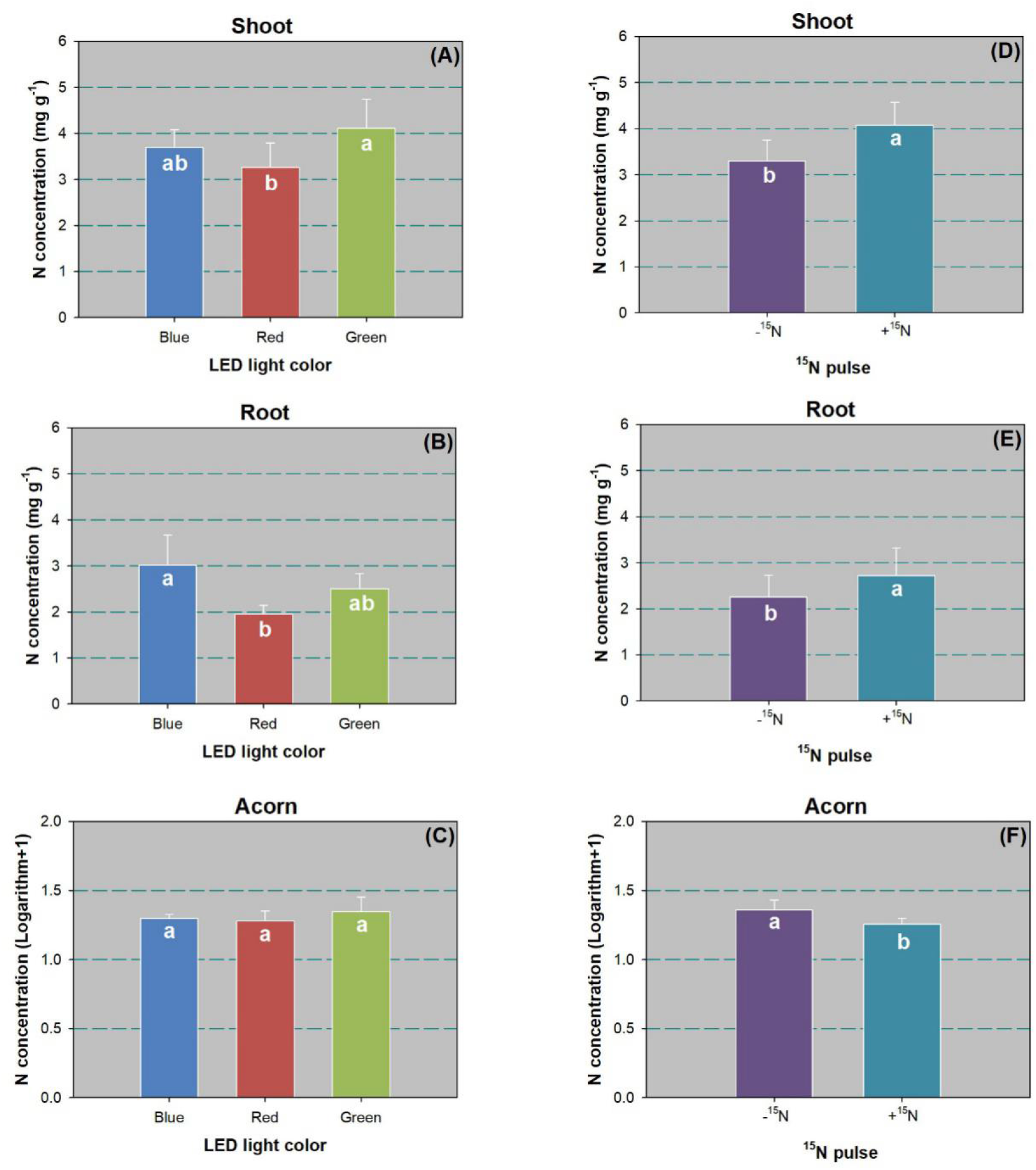
Nitrogen (N) concentration in shoot (A,D), root (B,E), and acorn (C,F) of Chinese cork oak (*Quercus variabilis* Blume) seedlings subjected to blue, red, and green colors of LED spectra (*n* = 24) or subjected to N pulse at 120 mg ^15^N plant^-1^ (+^15^N) or zero (-^15^N) (*n* = 36). Columns and bars represent means and standard errors, respectively. Different letters denote a significant difference according to Tukey’s test at the 0.05 level.

### N Content

Although the LED spectra had no effect on shoot N content, the ^15^N pulse treatment showed a significant effect (Table 3). Compared to the control, the ^15^N pulse increased shoot N content by 74% (Figure 6D). The effect of LED spectra on root N content was significant (Table 3). The red light spectrum resulted in lower root N content than the blue and green spectra (Figure 6B). Acorn N content was higher in the green light spectrum than in the blue light spectrum by 34% (Figure 6C). The ^15^N pulse resulted in a decline in acorn N content by 17% (Figure 6F).

**Figure 6.**
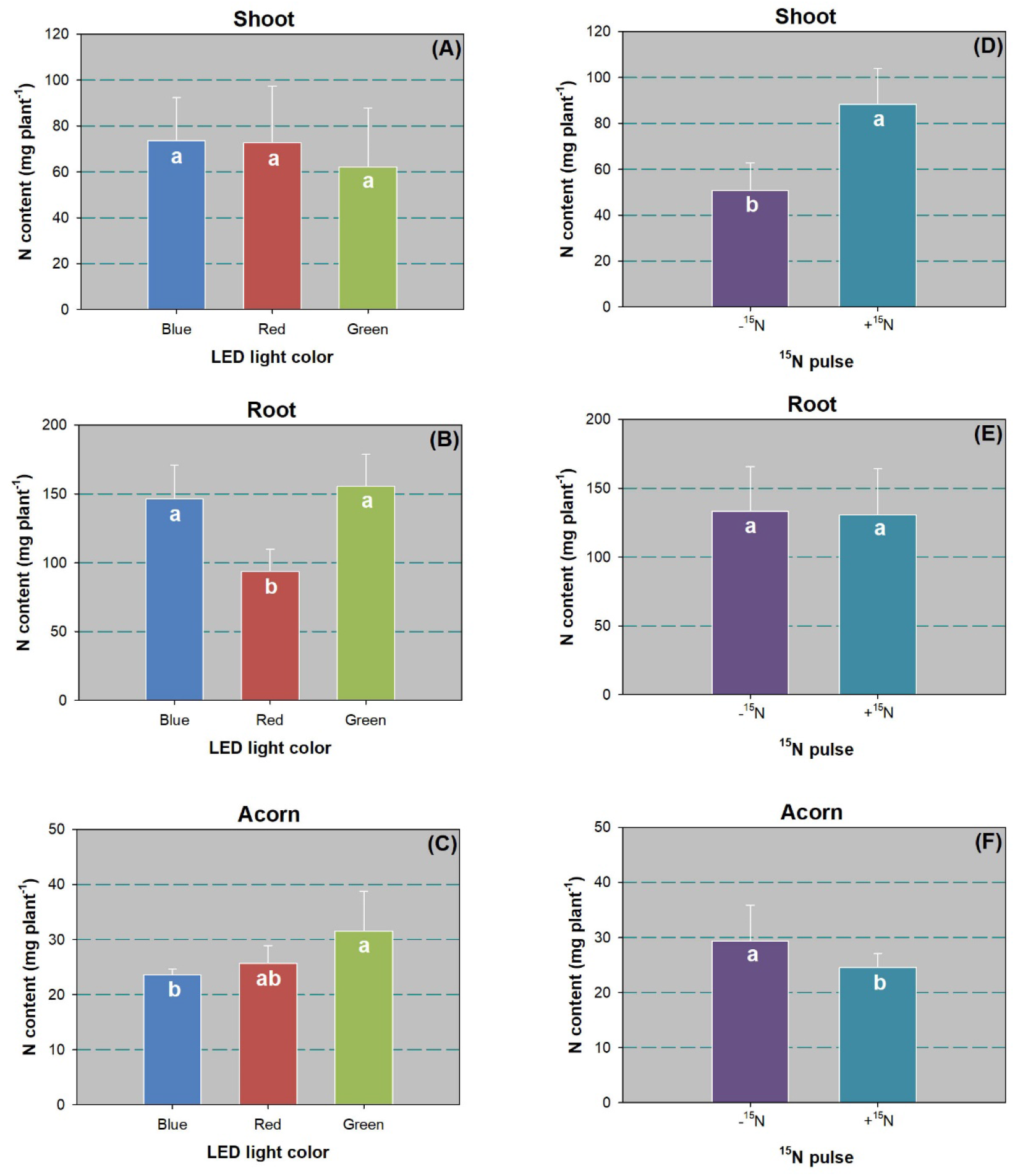
Nitrogen (N) content in shoots (A,D), roots (B,E), and acorns (C,F) of Chinese cork oak (*Quercus variabilis* Blume) seedlings subjected to blue, red, and green LED spectra (*n* = 24) or subjected to N pulse at 120 mg ^15^N plant^-1^ (+^15^N) or zero (-^15^N) (*n* = 36). Columns and bars represent means and standard errors, respectively. Different letters denote a significant difference according to Tukey’s test at the 0.05 level.

### Derived-N Percent and Amount

Although the LED spectra had no effect on the amounts of NDFP (*F*_2,6_ = 1.60, *P* = 0.2774) and NDFR (*F*_2,6_ = 0.47, *P* = 0.6482) in shoots (Figure 7A), their effects on derived-N percentage in shoots were significant for both NDFP (*F*_2,6_ = 7.75, *P* = 0.0217) and NDFR (*F*_2,6_ = 7.75, *P* = 0.0217). Both NDFP and NDFR percentages were higher in the blue light spectrum than in the red light spectrum (Figure 7D).

**Figure 7.**
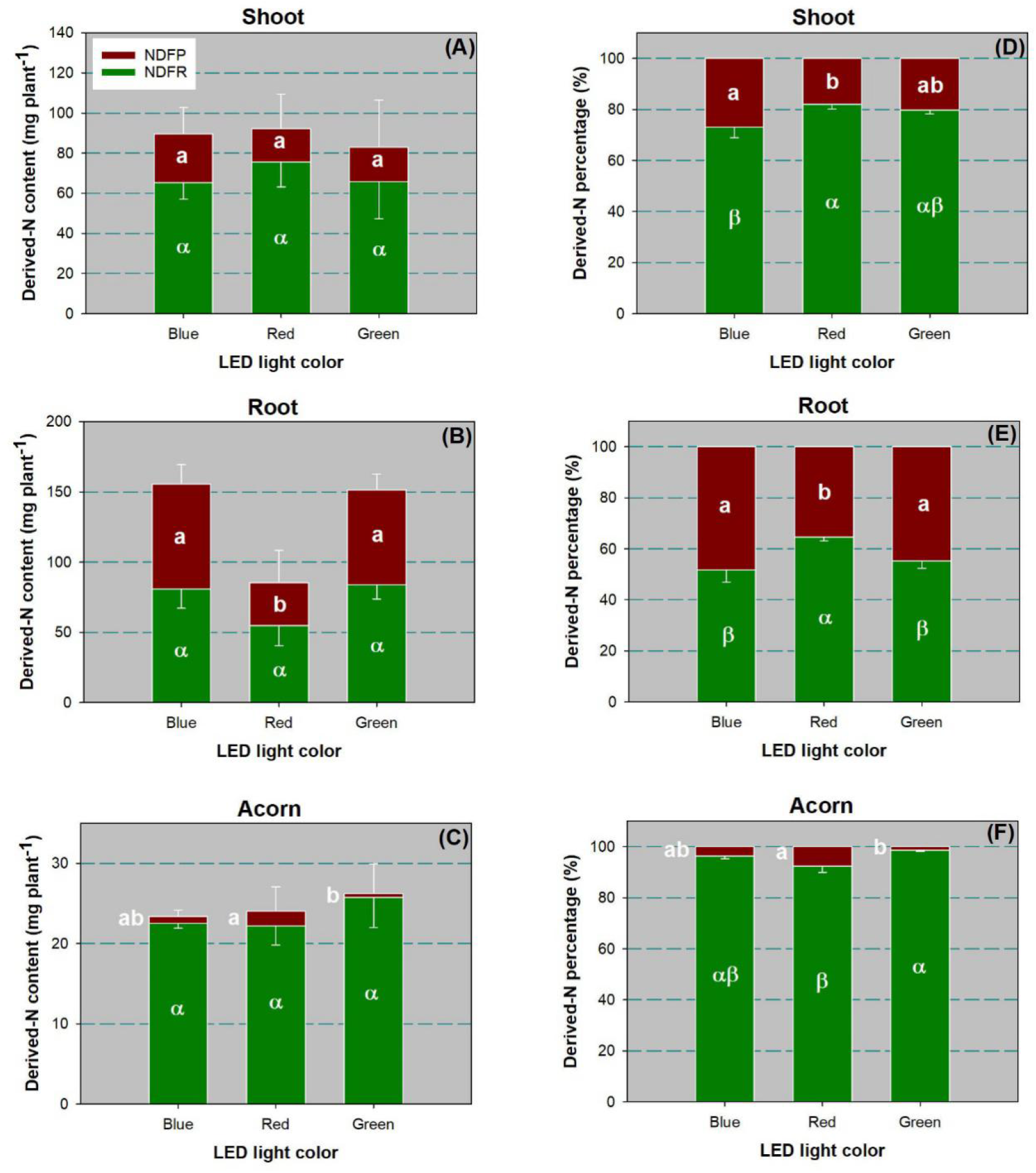
Amount and percent of nitrogen (N) derived from pulses (NDFP) and plant reserves (NDFR) in shoots (A,D), roots (B,E), and acorns (C,F) of Chinese cork oak (*Quercus variabilis* Blume) seedlings subjected to blue, red, and green LED spectra (*n* = 24). Columns and bars represent means and standard errors, respectively. Different letters denote a significant difference according to Tukey’s test at the 0.05 level.

The NDFP amount in roots was significantly affected by the LED spectra (*F*_2,6_ = 49.41, *P* = 0.0002). The red light spectrum resulted in a lower NDFP amount than the other two spectra (Figure 7B). In contrast, the percent of NDFR in roots was highest in the red light spectrum (*F*_2,6_ = 10.93, *P* = 0.0100), but the NDFP percentage was the lowest (*F*_2,6_ = 10.93, *P* = 0.0100; Figure 7E).

The NDFP amount in acorns was higher with the red light spectrum treatment than under green light (*F*_2,6_ = 10.93, *P* = 0.0100; Figure 7C). The percent of NDPF in acorns was higher in the red light spectrum treatment (*F*_2,6_ = 11.54, *P* = 0.0088; Figure 7F). In contrast, the acorn NDFR percentage was lower under the red light spectrum than under the green lighting (*F*_2,6_ = 11.54, *P* = 0.0088; Figure 7F).

## Discussion

We failed to find any significant effect of LED spectra on the growth of Chinese cork oak seedlings. These findings contradict those in the literature, but most previous studies about the growth of tree seedlings subjected to different LED spectra were not conducted with oak species. The current understanding about tree seedling response to LED spectra variation was mainly obtained from studies on coniferous species, such as Douglas fir (*Pseudotsuga menziesii*) [5], Engelmann spruce (*Picea engelmannii*) [5], Norway spruce (*Picea abies*) [15, 16], Scots pine (*Pinus sylvestris*) [15, 16], Prince Rupprecht’s larch (*Larix principis-rupprechtii* Mayr.) [10], Korean pine (*Pinus koraiensis*) [14], and Buddhist pine (*Podocarpus macrophyllus*) [12]. Oak seedling height showed rare variation when exposed to different LED spectra after a whole growing season [7, 8]. LED spectra also failed to show any significant effect on LA in *Q. ilex* seedlings [7], despite one report revealing a negative effect [49]. To the best of our knowledge, no study has investigated the RCD response to different spectra. However, a long-term field investigation revealed that across four upland oak species, RCD did not show any significant difference in sites with different understory transmittances until six years after seedling establishment [50]. As a hardwood species, Chinese cork oak seedlings may need longer and continuous investigation across years to detect significant changes in aerial organ growth.

Chinese cork oak seedlings also showed a null growth response to ^15^N pulses. This result surprised us because Uscola et al. [38] reported a significant growth response in *Q. ilex* to N input through exponential fertilization at a similar rate compared to the control. In another study with exponential fertilization of oak seedlings, growth of both *Q. rubra* and *Q. alba* seedlings highly increased with exposure to N addition at rates from 420 to 3350 mg N plant^-1^. Perhaps the N pulse used in our study was unable to induce the same growth response in oak seedlings as exponential fertilization. Exponential fertilization is a method of feeding tree seedlings in the nursery, which may follow a different pattern of growth than the natural N pulse. However, a two-year field investigation on simulated N deposition with Chinese cork oak saplings also found no shoot growth response [37], which strongly supports our findings. In another field study with long-term continuous N input, Yang et al. [51] found that growth of Konara oak (*Q. serrata*) seedlings did not significantly change in response to different N inputs until the third year of continuous fertilization. Again, like the response to LED spectra as concluded above, the growth of oak seedlings in response to N addition may need a study term of at least three years to record a significant response.

In leaves of Chinese cork oak seedlings, both chlorophyll and protein contents were higher in the blue light compared to the red light treatment. A similar response to blue and red spectra was also found in *Gerbera jamesonii* plantlets [52]. Li et al. [53] found higher foliar chlorophyll and protein contents in grape (*Vitis vinifera* L.) and further revealed the mechanism using RNA-seq analysis. They found that blue light can up-regulate genes that are related to microtubules, serine carboxypeptidase, and chlorophyll synthesis, and stimulate consumption of sugar to supply energy and down-regulated genes repressing protein for resistance. He et al. [54] obtained similar results with *Mesembryanthemum crystallinum* and found that low-wavelength blue light is necessary for chlorophyll synthesis through up-regulating N synthesis through promoting ribulose-1,5-bisphosphate carboxylase oxygenase (Rubisco) protein. This concurs with the GS activity results in our study and those of another study on *P. koraiensis* seedlings [14], which together suggest that the low-wavelength blue light can increase chlorophyll and protein contents through promoting N assimilation. This speculation is supported by our SLA results; SLA was lower under blue light than red light, suggesting a higher photosynthetic rate at the low cost of N consumption [55].

Biomass accumulation in either shoots or roots were significantly different between the blue and red spectra. This suggested that different spectra from blue and red lights had a similar impact on biomass production, although blue light resulted in a higher rate of N assimilation. Because we employed mixed wavelengths instead of monochromatic lights, the difference between our lights in blue and red colors was due to the change in the blue to red ratio. The lack of effect of different LED spectra with varied blue to red ratios on shoot and root biomass accumulation was also reported in studies on *Q. ilex* seedlings [8, 56] and coniferous seedlings, such as Engelmann spruce [5] and Norway spruce [16]. Thus, a wider range of tree and vegetable species may be used to demonstrate that different blue-to-red-light ratios produce different responses in shoot and root biomass [13, 14, 57]. Our findings, plus those of studies with similar results to ours, demonstrated the lack of biomass variation in oak seedlings when subjected to red and blue lights.

Chinese cork oak seedlings exposed to the green light spectrum had lower shoot biomass but more root biomass than the other two spectra, which resulted in a higher R/S. Most green light is reflected by the surface of aerial organs, which makes plants look green [58]. The low rate of green light photons by photosynthetic pigments limits the use of green light for photosynthetic production. This explains the reduced shoot biomass in the green light spectrum treatment and the promoted biomass allocation to roots. Our results are supported by a study on *Chrysanthemum morifolium* plants, where the higher R/S resulted from lowered shoot biomass but unchanged root biomass [59]. Another study used LED lighting on seven *Plectranthus scutellarioides* cultivars, similarly reporting lowered shoot biomass but increased R/S under green light exposure.

Notably, Chinese cork oak seedlings subject to the red light spectrum tended to show a lower level of N concentration in both shoots and roots, where those subjected to pulsed N had lowest tissue N pool percent. In addition, both the N content and NDFP amount in roots were the lowest in the red light treatment. In contrast, internal reserves of N derived from the plant accounted for the highest percent in the shoots and roots of seedlings subjected to the red light spectrum. Together, these results suggest that the red light spectrum restricted N uptake, resulting in a low competition ability for the internal reserves of N. Therefore, our first hypothesis is rejected. These results provide further explanation about low N concentration in shoots and roots is caused by weakened N uptake in the high red spectrum [10, 12, 14]. However, our findings, as well as some of those of other studies, do not agree with findings that N uptake in horticultural plants is promoted by red light [44, 60].

The declines in both N concentration and N content in acorns subjected to N pulse reflected the reduced reliance on remobilization from acorns to feed other issues when N availability is promoted [24]. Acorns did not show any responses in biomass or N concentration to LED spectra, which indicated that lighting the above-ground organs does not have any direct impact on biomass depletion and nutrient remobilization from acorns. However, the acorn was the only organ with higher NDFR under green light than under other lights, which led us to partly accept our second hypothesis. These results concur with those found in the field: certain abiotic factors can have significant effects on acorn quality [61, 62]. The higher N content in acorns treated with the green light spectrum resulted from the difference in the accumulative effect on errors in the ANOVA model of the product between biomass and concentration. Therefore, the third hypothesis is supported. The high N content in acorn can also be explained by the high percent of N derived from internal reserves in plants subjected to green light and the lesser reliance on N remobilization from acorns. The green light spectrum can promote root N uptake without heavy reliance on N derived from acorns, but the blue light spectrum showed a promotion of root N transported to shoots as demonstrated by the strong N assimilation response.

^15^N pulses can strongly increase N uptake and allocation to shoots, with promotion of chlorophyll content in leaves and N content in shoots. Other studies also reported that foliar N assimilation in trees can be increased by increasing ambient N deposition [63, 64]. We chose to feed oak seedlings with NH_4_-N as its assimilation with GS was sensitive to different spectra in *Fagus sylvatica* [49] and *P. koraiensis* seedlings [14]. As oak species prefer the uptake of NO_3_-N over NH_4_-N [65], our findings about the change in GS activities may be the result of the associated response of nitrate reductase (NR), which plays a key role in NO_3_-N assimilation. However, GS and NR showed a positive activity relationship with each other [14]; our results may have been at least partly the result of the NR driving force. The plant component uses of SMR and peat may have involved some microbiomes that affected the NH_4_-N and NO_3_-N balance relative to the pure substrates in sands. However, N assimilation was affected by the N amount trumping the mineral form, and our results regarding N uptake and assimilation were not likely affected by these basic conditions.

## 5. Conclusions

In this novel study, we tested the internal N recycling in oak seedlings subjected to different lighting spectra. A bioassay simulating N deposition in *Quercus variabilis* Blume under blue, red, and green spectra lighting provided by LED panels was conducted using seedlings subjected to ^15^N pulses. The blue light spectrum promoted higher percentage N in the roots, whereas the green light promoted N allocation to roots and strengthened the reliance on N remobilization from acorns. In contrast, the red light spectrum weakened N uptake and allocation relative to the blue light treatment. The ^15^N pulse promoted N uptake and assimilation but reduced N remobilization from acorns. Overall, in Chinese cork oak seedlings, the blue light spectrum benefits N use, the green light spectrum promotes N uptake, and the red light spectrum strengthens the remobilizing of N from acorns. N deposition did not show an interaction with light spectra but weakened the N remobilization from acorns without having a significant impact on growth and morphology. The ratio of red to far-red spectra has been proven to affect tree seedlings. Although we did not test far-red light in our spectra, further work is suggested to detect the response of internal N recycling of oak or other tree seedlings exposed to the far-red spectrum.

## Acknowledgments

Editors and reviewers who contribute to the improvement of this study. This research was funded by the Fundamental Research Funds for the Central Non-profit Research Institution of CAF (grant number CAFYBB2018ZB001).

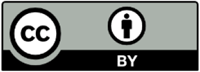 © 2020 by the authors. Submitted for possible open access publication under the terms and conditions of the Creative Commons Attribution (CC BY) license (http://creativecommons.org/licenses/by/4.0/).

## References

[1] McCreary DD, Tanaka Y, Lavender DP. Regulation of douglas-fir seedling growth and hardiness by controlling photoperiod. Forest Science. 1978; 24: 142–152.

[2] Arnott JT. Effect of light intensity during extended photoperiod on growth of amabilis fir, mountain hemlock, and white and Engelmann spruce seedlings. Canadian Journal of Forest Research. 1979; 9: 82–89.

[3] Bongarten BC, Hanover JW. Accelerating seedling growth through photoperiod extension for genetic testing: A case study with blue spruce (*Picea pungens*). Forest Science. 1985; 31: 631–643.

[4] Wei HX, Ren J, Zhou JH. Effect of exponential fertilization on growth and nutritional status in Buddhist pine (*Podocarpus macrophyllus* Thunb. D. Don) seedlings cultured in natural and prolonged photoperiods. Soil Sci Plant Nutr. 2013; 59: 933–941.

[5] Apostol KG, Dumroese RK, Pinto JR, Davis AS. Response of conifer species from three latitudinal populations to light spectra generated by light-emitting diodes and high-pressure sodium lamps. Canadian Journal of Forest Research. 2015; 45: 1711–1719.

[6] Zhu KY, Liu HC, Wei HX, Zhou JH, Zou QC, Ma GY, Zhang JQ. Prediction of nutrient leaching from culture of containerized Buddhist pine and Japanese maple seedlings exposed to extended photoperiod. Int J Agric Biol. 2016; 18: 425–434.

[7] Montagnoli A, Dumroese RK, Terzaghi M, Pinto JR, Fulgaro N, Scippa GS, Chiatante D. Tree seedling response to LED spectra: implications for forest restoration. Plant Biosyst. 2018; 152: 515–523.

[8] Smirnakou S, Ouzounis T, Radoglou KM. Continuous spectrum LEDs promote seedling quality traits and performance of *Quercus ithaburensis* var. *macrolepis*. Frontiers in Plant Science. 2017; 8.

[9] Li XW, Chen QX, Lei HQ, Wang JW, Yang S, Wei HX. Nutrient uptake and utilization by fragrant rosewood (*Dalbergia odorifera*) seedlings cultured with oligosaccharide addition under different lighting spectra. Forests. 2018; 9: 15.

[10] Zhao J, Chen X, Wei HX, Lv J, Chen C, Liu XY, Wen Q, Jia LM. Nutrient uptake and utilization in Prince Rupprecht’s larch (*Larix principis-rupprechtii* Mayr.) seedlings exposed to a combination of light-emitting diode spectra and exponential fertilization. Soil Sci Plant Nutr. 2019; 65: 358–368.

[11] Guo SL, Zhang S, Jia LW, Xu MY, Wang ZY. Root growth of Eleuthero (*Eleutherococcus senticosus* Rupr. & Maxim. Maxim.) seedlings cultured with chitosan oligosaccharide addition under different light spectra. Not Bot Horti Agrobot Cluj-Na. 2020; 48: 626–635.

[12] Luo YQ, Zhao SJ, Tang JY, Zhu H, Wei HX, Cui W, Wang MH, Guo P. White-light emitting diodes’ spectrum effect on photosynthesis and nutrient use efficiency in *Podocarpus macrophyllus* seedlings. J Plant Nutr. 2020: 9.

[13] Wei HX, Chen GS, Chen X, Zhao HT. Growth and nutrient uptake in *Aralia elata* seedlings exposed to exponential fertilization under different illumination spectra. Int J Agric Biol. 2020; 23: 644–652.

[14] Wei HX, Hauer RJ, Chen GS, Chen X, He XY. Growth, nutrient assimilation, and carbohydrate metabolism in Korean pine (*Pinus koraiensis*) seedlings in response to light spectra. Forests. 2020; 11: 18.

[15] Riikonen J. Pre-cultivation of Scots pine and Norway spruce transplant seedlings under four different light spectra did not affect their field performance. New For. 2016; 47: 607–619.

[16] Riikonen J, Kettunen N, Gritsevich M, Hakala T, Sarkka L, Tahvonen R. Growth and development of Norway spruce and Scots pine seedlings under different light spectra. Environ Exp Bot. 2016; 121: 112–120.

[17] Wei HX, Chen X, Chen GS, Zhao HT. Foliar nutrient and carbohydrate in *Aralia elata* can be modified by understory light quality in forests with different structures at Northeast China. Ann For Res. 2019; 62: 125–137.

[18] Giertych MM, Farrar JL. The effect of photoperiod and nitrogen on the growth and development of seedlings of Jack pine. Canadian Journal of Botany. 1961; 39: 1247–1257.

[19] Timmer VR. Exponential nutrient loading: A new fertilization technique to improve seedling performance on competitive sites. New For. 1997; 13: 279–299.

[20] Boivin JR, Salifu KF, Timmer VR. Late-season fertilization of *Picea mariana* seedlings: intensive loading and outplanting response on greenhouse bioassays. Ann For Sci. 2004; 61: 737–745.

[21] Oliet JA, Puertolas J, Planelles R, Jacobs DF. Nutrient loading of forest tree seedlings to promote stress resistance and field performance: a Mediterranean perspective. New For. 2013; 44: 649–669.

[22] Warren CR, Livingston NJ, Turpin DH. Response of douglas-fir seedlings to a brief pulse of N-15-labeled nutrients. Tree Physiol. 2003; 23: 1193–1200.

[23] Warren CR, Livingston NJ, Turpin DH. Photosynthetic responses and N allocation in Douglas-fir needles following a brief pulse of nutrients. Tree Physiol. 2004; 24: 601–608.

[24] Villar-Salvador P, Heredia N, Millard P. Remobilization of acorn nitrogen for seedling growth in holm oak (*Quercus ilex*), cultivated with contrasting nutrient availability. Tree Physiol. 2010; 30: 257–263.

[25] Salifu KF, Islam MA, Jacobs DF. Retranslocation, plant, and soil recovery of nitrogen-15 applied to bareroot black walnut seedlings. Commun Soil Sci Plant Anal. 2009; 40: 1408–1417.

[26] Du EZ, Terrer C, Pellegrini AFA, Ahlstrom A, van Lissa CJ, Zhao X, Xia N, Wu XH, Jackson RB. Global patterns of terrestrial nitrogen and phosphorus limitation. Nat Geosci. 2020; 13: 221–+.

[27] Kanakidou M, Myriokefalitakis S, Tsagkaraki M. Atmospheric inputs of nutrients to the Mediterranean Sea. Deep-Sea Res Part II-Top Stud Oceanogr. 2020; 171: 13.

[28] Gao Y, Zhou F, Ciais P, Miao CY, Yang T, Jia YL, Zhou XD, Klaus BB, Yang TT, Yu GR. Human activities aggravate nitrogen-deposition pollution to inland water over China. Natl Sci Rev. 2020; 7: 430–440.

[29] Wei H, He X. Foliar C/N stoichiometry in urban forest trees on a global scale. J For Res. 2020.

[30] Sun X, Kang H, Chen HYH, Bjorn B, Samuel BF, Liu C. Biogeographic patterns of nutrient resorption from Quercus variabilis Blume leaves across China. Plant Biol. 2016; 18: 505–513.

[31] Ma C, Zhang WH, Wu M, Xue YQ, Ma LW, Zhou JY. Effect of aboveground intervention on fine root mass, production, and turnover rate in a Chinese cork oak (*Quercus variabilis* Blume) forest. Plant Soil. 2013; 368: 201–214.

[32] Xin Y, Jia LY, Yuan JZ, Sun QS. A new cycloartane nortriterpenoid from *Quercus variabilis* Blume. Chin Chem Lett. 2009; 20: 817–819.

[33] Xu NN, Guo WH, Liu J, Du N, Wang RQ. Increased nitrogen deposition alleviated the adverse effects of drought stress on Quercus variabilis and Quercus mongolica seedlings. Acta Physiol Plant. 2015; 37: 11.

[34] Shi WH, Villar-Salvador P, Jacobs DF, Li GL, Jiang XX. Simulated predation of *Quercus variabilis* acorns impairs nutrient remobilization and seedling performance irrespective of soil fertility. Plant Soil. 2018; 423: 295–306.

[35] Wei HX, Zhao HT, Chen X, He XY. Secondary metabolites, carbohydrate accumulation, and nutrient uptake in *Aralia elata* (Miq.) Seem seedlings exposed to shoot cutting and different LED spectra. Acta Physiol Plant. 2020; 42: 15.

[36] Wei HX, Ma BQ, Hauer RJ, Liu CY, Chen X, He XY. Relationship between environmental factors and facial expressions of visitors during the urban forest experience. Urban Forestry & Urban Greening. 2020; 53: 10.

[37] Yu BY, Huang JG, Ma QQ, Guo XL, Liang HX, Zhang SK, Fu SL, Wan SQ, Yan JH, Zhang W. Comparison of the effects of canopy and understory nitrogen addition on xylem growth of two dominant species in a warm temperate forest, China. Dendrochronologia. 2019; 56: 8.

[38] Uscola M, Salifu KF, Oliet JA, Jacobs DF. An exponential fertilization dose-response model to promote restoration of the Mediterranean oak *Quercus ilex*. New For. 2015; 46: 795–812.

[39] Christian N, Herre EA, Clay K. Foliar endophytic fungi alter patterns of nitrogen uptake and distribution in *Theobroma cacao*. New Phytol. 2019; 222: 1573–1583.

[40] Warren CR, Livingston NJ, Turpin DH. Response of Douglas-fir seedlings to a brief pulse of ^15^N-labeled nutrients. Tree Physiol. 2003; 23: 1193–1200.

[41] James JJ, Richards JH. Plant nitrogen capture in pulse-driven systems: interactions between root responses and soil processes. J Ecol. 2006; 94: 765–777.

[42] Xu L, Zhang X, Zhang DH, Wei HX, Guo J. Using morphological attributes for the fast assessment of nutritional responses of Buddhist pine (*Podocarpus macrophyllus* Thunb. D. Don) seedlings to exponential fertilization. PLoS One. 2019; 14: 14.

[43] Zhu H, Zhao SJ, Yang JM, Meng LQ, Luo YQ, Hong B, Cui W, Wang MH, Liu WC. Growth, nutrient uptake, and foliar gas exchange in pepper cultured with un-composted fresh spent mushroom residue. Not Bot Horti Agrobot Cluj-Na. 2019; 47: 227–236.

[44] Wang R, Wang Y, Su Y, Tan JH, Luo XT, Li JY, He Q. Spectral effect on growth, dry mass, physiology and nutrition in *Bletilla striata* seedlings: Individual changes and collaborated response. Int J Agric Biol. 2020; 24: 125–132.

[45] Wei HX, Zhao HT, Chen X. Foliar N:P stoichiometry in *Aralia elata* distributed on different slope degrees. Not Bot Horti Agrobot Cluj-Na. 2019; 47: 887–895.

[46] El Zein R, Bréda N, Gérant D, Zeller B, Maillard P. Nitrogen sources for current-year shoot growth in 50-year-old sessile oak trees: an in situ ^15^N labeling approach. Tree Physiol. 2011; 31: 1390–1400.

[47] Wei HX, Xu CY, Ma LY, Wang WJ, Duan J, Jiang LN. Short-term nitrogen (N)-retranslocation within *Larix olgensis* seedlings is driven to increase by N-deposition: Evidence from a simulated N-15 experiment in Northeast China. Int J Agric Biol. 2014; 16: 1031–1040.

[48] Liu ZL, Hikosaka K, Li FR, Jin GZ. Variations in leaf economics spectrum traits for an evergreen coniferous species: Tree size dominates over environment factors. Funct Ecol. 2020; 34: 458–467.

[49] Astolfi S, Marianello C, Grego S, Bellarosa R. Preliminary investigation of LED lighting as growth light for seedlings from different tree species in growth chambers. Not Bot Horti Agrobot Cluj-Na. 2012; 40: 31–38.

[50] Brose PH, Rebbeck J. A comparison of the survival and development of the seedlings of four upland oak species grown in four different understory light environments. J For. 2017; 115: 159–166.

[51] Yang AR, Hwang J, Cho MS, Son Y. The effect of fertilization on early growth of konara oak and Japanese zelkova seedlings planted in a harvested pitch pine plantation. J For Res. 2016; 27: 863–870.

[52] Meng XY, Wang Z, He SL, Shi LY, Song YL, Lou XY, He D. LED-supplied red and blue light alters the growth, antioxidant status, and photochemical potential of in vitro-grown *Gerbera jamesonii* plantlets. Hortic Sci Technol. 2019; 37: 473–489.

[53] Li CX, Xu ZG, Dong RQ, Chang SX, Wang LZ, Khalil-Ur-Rehman M, Tao JM. An RNA-seq analysis of grape plantlets grown in vitro reveals different responses to blue, green, red LED light, and white fluorescent light. Frontiers in Plant Science. 2017; 8: 16.

[54] He J, Qin L, Chong ELC, Choong TW, Lee SK. Plant growth and photosynthetic characteristics of *Mesembryanthemum crystallinum* grown aeroponically under different blue-and red-LEDs. Frontiers in Plant Science. 2017; 8: 13.

[55] El-Serafy RS. Phenotypic plasticity, biomass allocation, and biochemical analysis of cordyline seedlings in response to oligo-chitosan foliar spray. J Soil Sci Plant Nutr. 2020: 12.

[56] Zhang T, Shi YY, Piao FZ, Sun ZQ. Effects of different LED sources on the growth and nitrogen metabolism of lettuce. Plant Cell Tissue Organ Cult. 2018; 134: 231–240.

[57] Fraszczak B. Effect of short-term exposure to red and blue light on dill plants growth. Hortic Sci. 2013; 40: 177–185.

[58] Folta KM, Maruhnich SA. Green light: a signal to slow down or stop. J Exp Bot. 2007; 58: 3099–3111.

[59] Jeong SW, Park S, Jin JS, Seo ON, Kim GS, Kim YH, Bae H, Lee G, Kim ST, Lee WS, Shin SC. Influences of four different light-emitting diode lights on flowering and polyphenol variations in the Leaves of Chrysanthemum (*Chrysanthemum morifolium*). J Agric Food Chem. 2012; 60: 9793–9800.

[60] Heo JW, Lee YB, Kim DE, Chang YS, Chun C. Effects of supplementary LED Lighting on growth and biochemical parametets in *Dieffenbachia amoena* ‘Camella’ and *Ficus elastica* ‘Melany’. Korean J Hortic Sci Technol. 2010; 28: 51–58.

[61] Bogdziewicz M, Crone EE, Steele MA, Zwolak R. Effects of nitrogen deposition on reproduction in a masting tree: benefits of higher seed production are trumped by negative biotic interactions. J Ecol. 2017; 105: 310–320.

[62] Brooke JM, Basinger PS, Birckhead JL, Lashley MA, McCord JM, Nanney JS, Harper CA. Effects of fertilization and crown release on white oak (*Quercus alba*) masting and acorn quality. For Ecol Manage. 2019; 433: 305–312.

[63] Serapiglia MJ, Minocha R, Minocha SC. Changes in polyamines, inorganic ions and glutamine synthetase activity in response to nitrogen availability and form in red spruce (*Picea rubens*). Tree Physiol. 2008; 28: 1793–1803.

[64] Liu N, Wu SH, Guo QF, Wang JX, Cao C, Wang J. Leaf nitrogen assimilation and partitioning differ among subtropical forest plants in response to canopy addition of nitrogen treatments. Sci Total Environ. 2018; 637: 1026–1034.

[65] Truax B, Lambert F, Gagnon D, Chevrier N. Nitrate reductase and glutamine synthetase activities in relation to growth and nitrogen assimilation in red oak and red ash seedlings: effects of N-forms, N concentration and light intensity. Trees-Struct Funct. 1994; 9: 12–18.

